# Tetrahydrocannabinol exposure to postejaculatory sperm compromises sperm structure, function, the epigenome, and early embryo development

**DOI:** 10.64898/2026.03.23.713385

**Authors:** Muhammad S. Siddique, Santosh Anand, João D de Agostini Losano, Zongliang Jiang, Ramji K. Bhandhari, Bradford W. Daigneault

## Abstract

Cannabis (marijuana) is the most widely used recreational drug in the USA accounting for about 62 million users in 2024. Among cannabis users, 26% are of prime reproductive age (18-25 years). Delta-9 tetrahydrocannabinol (THC) is the principal psychoactive component of cannabis and has been detected in human seminal fluids. Although abundant evidence indicates adverse effects of THC exposure on spermatogenesis in different species, acute effects of THC on postejaculatory sperm including fertilization potential and subsequent carryover effects on embryo development are largely unknown. The present study was designed to provide missing information on structural and mechanistic effects of THC exposure to postejaculatory sperm function by evaluating sperm indices often overlooked or masked during clinical evaluation. A bovine embryo continuum model was employed to determine effects of THC on sperm structure, kinematics, bioenergetics, and binding mechanisms. Effects of THC on the sperm genomic and epigenomic landscape were determined, complemented by paternal carry over effects on embryo development as a human translational model to elucidate paternal effects on future development, and to mirror sperm exposure during transport within the female reproductive tract. Cryopreserved bovine sperm from three bulls were independently exposed to physiologically relevant concentrations of THC (0 and 32nM, n = 2 individual replicates/bull) for 24 h under non-capacitating conditions at 25°C followed by quantification of sperm kinematics at 37°C. Samples of THC-exposed sperm and vehicle-control (0.1% DMSO) were collected in replicate following immediate addition of THC (0 h) and again at 24 h. DNA damage, acrosome integrity, bioenergetics, changes to DNA methylation and embryo development were quantified. Data were analyzed by logistic regression with a generalized linear mixed effect model. Computer-assisted sperm assessment revealed a reduction in progressive motility of THC-exposed sperm after 24 h while other parameters were not affected. Acrosome integrity as determined by flowcytometric analysis with FITC-PSA was severely compromised in THC-exposed sperm (P ≤ 0.05), despite no detectable difference in capacitation status using merocyanine staining. Similarly, DNA integrity as determined by TUNEL assay was significantly impaired after 24 h of THC exposure (P ≤ 0.05). Mechanistic effects of THC were explored through characterization of the transmembrane G-protein coupled cannabinoid 1 receptor (CB1). CB1 is expressed in the post-acrosomal region and its abundance decreased as compared to unexposed sperm. Alterations to the methylation landscape of sperm were then determined after 24 h of THC exposure through whole-genome Enzymatic Methyl Sequencing. PCA analysis indicated that sperm from different males formed distinct clusters, implying individual differences among bulls, while the effects of THC exposure produced tighter clusters. Paternal carryover effects on embryos derived by in vitro fertilization from THC exposed sperm had reduced 2-cell cleavage, 8-16 cell morula development, and reduced blastocyst development compared to unexposed sperm (46% vs. 33%). In conclusion, post-ejaculatory mammalian sperm exposure to THC compromises acrosome integrity, induces DNA damage, changes the sperm methylome, and reduces developmental potential. Collectively, these data implicate new considerations for recreational and clinical use of cannabis that impact cellular and molecular mechanisms important for sperm function with detrimental consequences for gamete interaction and embryo development.

## Introduction

Cannabis (marijuana) is the most used recreational drugs in the USA accounting for about 62 million users (18% of total population) in 2024 [1]. Similarly, a staggering 75% increase from 2010-2022 in global cannabis users of reproductive age was reported to reach 228 million[2]. In the USA, around 42% of young people from 19-30 years of age reported using cannabis in 2023 [3]. Additionally, there is a four-fold increase in cannabis use in the USA from 2016 to 2020 for various medical conditions [4]. The increased social acceptance of cannabis use in people of prime reproductive age is concerning due to unknown effects on early embryo development. Although cannabis is associated with reproductive disorders that affect spermatogenesis, steroidogenesis, and negatively regulate seminal parameters including volume, motility, viability and morphology [5–12], there are few comprehensive studies to determine the relationship between cannabis use and male factor infertility despite chronic use in young people.

Delta-9-tetrahydrocannabinol (THC) is the main psychoactive chemical in cannabis [13–15]. An abundance of data exists to describe negative effects of THC on sperm at the testicular level in different species [16–24], but a large gap of knowledge remains to describe influences of THC on postejaculatory sperm function following acute exposure from short-term use prior to gamete interaction [25, 26]. THC acts as a partial agonist at transmembrane G-protein coupled cannabinoid receptors 1 and 2 (CB1/2), included in the endogenous endocannabinoids system (ECS). Binding of THC with these receptors can initiate downstream intracellular events in other cell types, but these mechanisms have not been explored in mature sperm [27, 28]. The ECS contains endogenously produced ligands such as anandamide (AEA) and 2-arachidonylglycerol (2-Ag) coined as endocannabinoids (eCBS) [29, 30]. ECS is involved with essential male reproductive function of spermatogenesis, steroidogenesis, and sperm function [31–33]. Observational studies indicate that stimulation of the ECS and reduced AEA levels negatively affect sperm motility and viability in humans while in boar sperm it alters capacitation and affects acrosome integrity [15, 27–29].

THC and its primary metabolites (11-hydroxy-THC and 11-nor-9-carboxy-THC) have been detected and quantified in blood and seminal plasma of chronic cannabis users with a mean value of 10.1ng/mL and 71.6ng/mL, respectively [34]. Concentration of the THC metabolites in blood plasma rises with increasing Δ9-THC concentration implying a relationship between concentration of THC and its metabolites [34, 35]. Major exposure routes of postejaculatory sperm to THC include seminal plasma and through fluids encountered during transport within the female reproductive tract [36, 37]. Effects of medical and recreational exposure of THC on human sperm include a decrease in progressive motility, inhibition of spontaneous acrosome reaction, changes in the epigenetic landscape, and affected behavioral endpoints in offspring [38–43]. Although these effects likely disrupt sperm function, direct consequences and mechanisms of action have not been fully explored. Furthermore, mechanisms uncoupling testicular exposure and transient postejaculatory sperm exposure are lacking. Mammalian postejaculatory spermatozoa are specialized transcriptionally quiescent cells that contain highly compacted chromatin due to histones replacement by protamines during spermatogenesis. This fundamental change in sperm chromatin structure likely provides protection from environmental stressors during sperm transport within the female reproductive tract. Although highly condensed DNA of sperm yields heterochromatin structures, approximately 8% of the human and bovine sperm chromatin structure remain open (euchromatin), lending susceptibility to THC exposure from cannabis [44–47]. A wide range of environmental factors including endocrine disrupting chemicals, drugs, radiations, lifestyles choices and diet can induce epigenomic modifications. These modifications can influence phenotypic outcomes including disease processes, and, when occurring in the germline, facilitate epigenetic inheritance [48, 49]. The sperm epigenome contains important information that although likely inconsequential for sperm function prior to fertilization, plays critical regulatory zygotic roles that influence early embryo development [50]. Changes to the sperm epigenome may lead to transcription alteration of genes involved in early embryo development that have a profound effect on neural system development, cancer cell proliferation, imprinted regions, and subsequently affect generational inheritance [51–54]. Aberrant sperm DNA methylation is associated with imprinting disorders and subfertility with mechanisms attributed to aberrant transcription and changes in methylation levels of differentially methylated regions (DMRs) necessary for fertility that are involved in spermatogenesis and early embryonic development [53, 55].

In rodents, chronic THC paternal exposure both as a sub cutaneous injection and oral gavage prior to mating causes methylation changes in rat sperm, long-term behavioral differences, neurological disorders and impairs cognitive function in offspring when mated to naïve females [56–61]. Similarly, sperm intrinsic factors such as coding and non-coding RNAs (miRNAs) are transferred during fertilization and are important epigenetic contributors and post transcriptional modulators for early embryonic development [62–64]. These miRNAs which can serve as fertility biomarkers for in vitro fertilization (IVF) are potential targets of THC insult, but direct mechanisms have not been explored [62, 65, 66]. Furthermore, unlike somatic cells, postejaculatory sperm are transcriptional silent, thereby confounding genomic and epigenomic repair mechanisms and thus likely elevating consequences of impacts to sperm.

The present study was aimed at determining the structural, genomic, and epigenomic effects of THC exposure to postejaculatory sperm and carryover effects on the embryo. Studies were further aimed at characterization of novel epigenetic mechanisms that directly impact sperm function and subsequent embryo development to reveal developmental deficiencies of paternal origin.

## Methods

Experiments conducted herein utilized commercially available frozen-thawed bull sperm from ORIgen (Huntley, MT, USA). Live animals were not used for these experiments. Unless otherwise stated, all reagents were purchased from Sigma Aldrich^®^ (St. Louis, MO, USA).

### Isolation of bovine sperm

Unsorted frozen-thawed sperm from three different commercial bulls of proven reproductive performance were used for all experiments unless stated otherwise. In each replicate, four frozen straws (0.5mL) were thawed at 37^°^C for 30s followed by isolation on a discontinuous Percoll gradient (90:45%) and washed twice by centrifugation in HEPES buffered saline in accordance with established protocols. Sperm pellets were then resuspended to a final concentration of 20 ×10^6^ total sperm/mL in 400uL volumes of a non-capacitating Tyrode’s albumin lactate pyruvate (TALP) medium free of bicarbonate and albumin (99 mM NaCl, 24.8mM NaHCO_3_,10mM HEPES, 0.33mM NaH_2_PO_4_, 24mM sodium lactate (60%), 2.4mM MgCl_2_ · 2H_2_O, 2.6 mM CaCl_2_ · 2H_2_O [67].

### Preparation of *THC* for sperm exposure

A stock solution of THC (320uM) was prepared from initial stock of 3.17mM (Millipore Sigma Cat No: 1651621) in 100% dimethyl sulfoxide (DMSO). From the 320uM stock solution, an additional stock of 32uM solution was prepared using 100% DMSO and stored in glass scintillating vials wrapped in paraffin at room temperature. On the day of experiment, 10uL of 32uM THC was added to 90uL of TALP medium to obtain 3.2uM (3200nM) working dilutions of final THC concentration in 10% DMSO. Then, 1:100 final dilutions in TALP media containing sperm were made to achieve final THC concentrations of 32.0nM in 0.1% DMSO including a vehicle control (VC - DMSO 0.1%) prepared in the same manner but devoid of THC. The THC concentration used in the experiments is reflective of a range that includes plasma THC levels achieved after recreational cannabis use in humans [25, 26, 68–73] and also quantification of THC in human seminal plasma reported in chronic cannabis users [35].

### Sperm exposure to *THC*

Sperm samples were maintained at a final concentration of 20×10^6^ total sperm/mL in TALP and were exposed to a single concentration of THC (32nM), simultaneously with VC samples (0.1% DMSO). All samples were maintained for up to 24 h at ambient (25^°^C) temperature to replicate temporal aspects of sperm transport within the female reproductive tract. An aliquot of each sample was warmed to 37°C for 10 min prior to sperm kinematic analyses.

### Detection and characterization of cannabinoid recptor-1 (CB1)

Characterization of CB1 abundance was performed with THC-exposed and unexposed sperm by immunocytochemistry (ICC) and western blot (WB) analyses. All washes were performed 3 times at room temperature with PBS for ICC and with TRIS Buffered Saline containing 0.1% Tween **^®^** 20 (TBST) for WB. For ICC, 50µL of sperm extended in TALP at 20 ×10^6^ / total sperm/mL was placed on a glass slide and dried at room temperature followed by fixation with 4% paraformaldehyde for 30 min. Fixed sperm were then washed and permeabilized with TritonX 0.1% for 15 min at room temperature and then washed again. Each slide was then incubated with 200µL of Blocking Buffer (BB) for 2h at room temperature and then washed. Primary antibody incubation for detection of CB1 include the addition of 50 µL of Anti-CB1(proteintech^®^ Cat No: 17978-1-AP) at a dilution of 1:50 in BB for 15 h at 4^°^C. Slides were then washed and 100µL of secondary antibody (Donkey-AntiRabbit Abcam Cat No: ab6799) was added at a dilution of 1:100 (in BB) for 1 h in the dark followed by washing. Finally, 10µL of Vector mounting medium with DAPI (#H1200, Vector^®^) was added followed by placement of coverslip. Slides were either analyzed immediately or stored at 4^°^C for future analysis. Fluorescence emission and detection were performed at wavelengths of 360/432 (DAPI), and 541/595 (Rhodamine), respectively. Emission was detected with an AURAIII-UCGRnIR Light Engine (LED Lumencor 4-channel) compatible with Penta band filter cube on a Nikon TE2000U inverted microscope at 200X magnification and Nikon Elements software. A similar procedure was adopted for human and bovine placenta and mouse brain, serving as positive controls for CB1 detection in these experiments.

For WB analysis, CB1 detection was performed following isolation of frozen-thawed bull sperm as described above. Proteins were extracted by diluting the sperm pellet (100 µL) after centrifugation (500×G for 10 min) in 100 µL of RIPA buffer containing dithiothreitol (DTT)(Cat. #10708984001, Sigma^®^), phosphatase inhibitor (Cat. #78420, Thermo Fisher^®^), and protease inhibitor (Cat. # 11836170001, Sigma^®^), at a 2:1 (v:v) dilution. Samples were kept on ice for 30 min and vortexed every 10 min before storage at −80°C. On the day of the assay, protein from sperm lysates (100 µg) was solubilized in 4X Laemmli buffer (Cat. # 1610747, BioRad^®^) by dilution to 1X in sample with 10% β-mercaptoethanol, heated at 95°C for 10 min and resolved by electrophoresis gel in 8-10% gels (Mini-PROTEAN^®^ TGX^™^, Bio-Rad, Hercules, CA) for 1.5 h at 115 V. The resolved proteins were transferred to a nitrocellulose membrane in transfer buffer (48 mM Tris base, 139 mM glycine, 20% methanol, and 0.025% SDS) for 1 h at 95 V. The membrane was blocked at room temperature in tris-buffered saline, 0.1% Tween (TBST) buffer (20 mM Tris Base, 137 mM NaCl, 0.1% Tween20 and 5% powdered milk) and washed. Protein detection was performed with Anti-CB1(1:1500 proteintech Cat No: 17978-1-AP) in TBST buffer overnight at 4°C followed by detection with anti-rabbit secondary antibody HRP-linked (1:2000; Cat. #7074P2, Cell Signaling Technology^®^), in TBST buffer. After washing, bound antibody was visualized by BIO-RAD ChemiDoc XRS Imaging System and Software Image Lab (BIO-RAD). Positive (equine pituitary) and negative controls (RIPA extraction buffer) were utilized to ensure specificity and authenticity of single-band detection. Antibody specificity was further validated by probing with secondary antibody only.

### Sperm kinematic assessment

Sperm kinematic parameters were objectively assessed by computer assisted sperm analysis (CASA-IVOS System, Hamilton Thorne, Beverly, MA) [74]. For each experiment, 2 replicates per bull were performed. Time points for kinematic assessment included 0 h (within 5 -10 minutes of THC exposure), and again at 24 h following maintenance at ambient temperature as previously described and then warming of an aliquot to 37°C for 10 min prior to all analyses. Parameters acquired include total motility (TM), progressive motility (PM), average path velocity (VAP) and curvilinear velocity (VCL). A minimum of five fields and a total of 500 spermatozoa were counted on a heated stage at 37°C by acquiring 30 frames at a rate of 60 Hz with a minimum contrast of 15 and minimum cell size of 10 pixels. Progressive minimum cutoff values included VAP at 20 µm/S, slow cell cut off VAP at 5 µm/S, and VCL cut off is 6 µm/S. All CASA measurements were acquired using Leja 4 chamber 20µM slides with the addition of 3µL of sample in each chamber. (IMV, 0251070).

### DNA integrity measurement by terminal deoxynucleotidyl transferase *dUTP nick end labeling (TUNEL ASSAY)*

Terminal deoxynucleotidyl transferase dUTP nick end labeling (TUNEL) was used to quantify DNA integrity following sperm exposure to THC using a One-step TUNEL In Situ Apoptosis Kit (Green-FITC) (Elabscience, Cat No. E-CK-A320). Sensitivity and specificity of the kit were determined with the use of controls consisting of no enzyme (negative) and the addition of DNASe (positive) to ensure ≥85% labelling in exposed sperm. In this assay, 50uL of sperm samples previously maintained at a concentration of 20×10^6^ total sperm /mL were placed on a glass slide at 37^°^C for air drying. Paraformaldehyde (4%) was then added to each sample in 100µL volumes for fixation at a room temperature for 30 min followed by PBS washing. Sperm were then permeabilized by incubation with TritonX 0.1% for 15 minutes at 37^°^C followed by PBS washing. After washing, 100µL of Terminal deoxynucleotidyl transferase (TdT) equilibration buffer was added to each slide and incubated for 30 minutes at 37^°^C followed by addition of 50 µL of labelling solution (TdT Equilibration buffer, labeling solution and TdT enzymes) and incubation again at 37^°^C for 4 h in a humidified chamber without light. Slides were then washed with PBS and 100µL of DAPI working solution (25µg/mL) was added for 15 minutes at room temperature followed by a PBS wash. Slides were air dried and analyzed immediately or held at 4^°^C for future analysis. Two replicates for each bull (3 bulls) were performed with a minimum of 120 sperm per slide enumerated per replicate for a total of 960 sperm for each treatment. Fluorescence emission and detection were performed at wavelengths of 360/432 (DAPI), and 461/515 nm (FITC), respectively. Emission was detected with an AURAIII-UCGRnIR Light Engine (LED Lumencor 4-channel) compatible with Penta band filter cube on a Nikon TE200U inverted microscope at 200X magnification and Nikon Elements software.

### Detection and quantification of acrosome integrity and capacitation status by flow cytometry

Acrosome detection and capacitation status of sperm were quantified by flow cytometry following sperm isolation and THC exposure as described above. Four Fluorochromes were used for all analyses. FITC-PSA was used to determine acrosome integrity. Hoechst (H33342) and propidium iodide (PI) were used to calculate membrane integrity and live dead ratio, and merocyanine (M540) probe confirmed capacitation status of sperm. Prior to analyses, fluorochrome minus one (FMO) was utilized as a negative control for each fluorochrome. For each fluorochrome, an FMO control was established by adding the other three fluorochromes to the sperm sample and omitting the one tested. This methodology produced four separate FMOs (each for one particular fluorochrome) and then positive and negative signals were plotted to confirm the absence of fluorochrome that was excluded in that particular FMO. Positive controls for viability included sperm samples held at 95°C to validate >90% dead sperm detected by PI positive matrix. For actual analysis of THC exposed and unexposed sperm, frozen-thawed bovine sperm from three bulls were isolated on 90:45 Percoll gradient, pooled and brought to a final concentration to 20×10^6^ total sperm/mL in TALP. Aliquots (500µL) of sperm from THC exposed and unexposed samples were included for final analysis with the addition of FITC-PSA (100ug/mL), H342 (5mg/mL), PI (1mg/mL) and Merocyanine540 (M540 100uM) for 15 min incubation at 37°C in dark. Sperm samples with incubated fluorochrome were then evaluated by flow cytometry using a Thermofisher Attune Nxt Flow Cytometer as described [75]. Gating strategy includes the following: After gating live sperm, capacitated sperm were detected as merocyanine+ and acrosome reacted cells as FITC-PSA+. Compensation was performed by using sperm samples to obtain a positive and negative population for each fluorochrome and the compensation matrix was plotted using FlowJo software (Version 10.0.7, Treestar, Palo Alto, CA). FITC-PSA was detected at 494/520 nM using BL1 filter while PI was detected at 537/618 nM using BL3 filter, H342 was detected at 405/454 nM by VL1 and M540 was detected at 560/579nM using a BL2 filter. For acrosome and plasma membrane integrity of THC-exposed and unexposed control sperm, FITC-PSA was plotted against PI to analyze acrosome reacted and non-acrosome reacted while also determining membrane integrity simultaneously. FITC-PSA-/PI- staining represents the non-acrosome reacted viable sperm population, FITC-PSA+/PI- staining indicates acrosome reacted viable sperm population, FITC-PSA+/PI+ positive indicates acrosome reacted non-viable sperms, and FITC-PSA-/PI+ non-acrosome reacted non-viable sperms. To analyze capacitation level of THC exposed sperm, merocyanine (M540) was plotted against PI. The gating strategy was similar as described above. M540-/PI- staining represents the non-capacitated viable sperm population, M540+/PI- staining indicates capacitated viable sperm population, M540+/PI+ positive indicates capacitated non-viable sperms, and M540-/PI+ non-capacitated non-viable sperms. All Data were analyzed using FlowJo software (Version 10.0.7, Treestar, Palo Alto, CA). Once data tables were extracted from FlowJo, averages between the 2 panels were calculated. The flow cytometer settings were configured to collect a volume of 200 µL of sample or a maximum of 100,000 cells, whichever was reached first.

### High-resolution respirometry analysis of THC exposed sperm

The Oxygraph-2k (Oroboros Instruments, Innsbruck, Austria) was used for measuring sperm oxygen flux, electron transport (ET) capacity, and OXPHOS capacity in 0.5 mL chambers as previously optimized for bull sperm [76, 77]. Before starting the assay, calibration at air saturation versus zero oxygen was performed by allowing the media (0.5 mL of TALPNC) to equilibrate with air in the oxygraph closed chambers and stirred at 540–560 rpm for 30–40 min until a stable signal was detected. After calibration, the chambers were opened, 300 µL of media was removed and 300 µL of sperm at 20 x10^6^/total sperm mL (6 x 10^6^ total sperm) of both treatment (THC) and VC was added to each chamber (A and B). To minimize intra-assay variability, sperm samples from treatment and unexposed control were used in the same assay, alternating the groups across chambers (A and B) for each assay. The respirometry assay was initiated by first detecting basal respiration. After oxygen consumption signals stabilized, reagents targeting independent components of the mitochondrial electron transport chain (ETC) were injected sequentially using equipment-specific glass syringes, following the same principle. Reagents were injected in the following steps: 1) Sperm addition (no chemical), 2) Oligomycin 5nM (ATP synthase inhibitor), 3) ADP 2.5 mM + Succninate 10mM (mitochonadrial stimulation), 4) CCCP 0.5 µM (Carbonyl cyanide m-chlorophenylhydrazone, mitocondrial uncoupler), 5) Rotenone 0.5 µM (complex I inhibitor) and 6) Antimycin 2.5 µM (complex III inhibitor). From these steps, the following respirometric variables were generated after the addition of sperm: 1) Basal respiration = Rotenone + Antimycin injection (non-mitochondrial respiration), 2) ATP-linked respiration = after sperm added - aftter oligomycin injection, 3) Proton leak = after oligomycin injection - Rotenone + Antimycin injection (non-mitochondrial respiration), 4) ADP-dependent = after ADP injection - after oligomycin injection, 5) Spare respiratory capacity = after CCCP injection - basal respiration, 6) Maximal respiration = after CCCP injection - Rotenone + Antimycin injection (non-mitochondrial respiration), 7) Complex I-dependent= after CCCP injection - after rotenone injection, 8) Complex III-dependent = after Rotenone injection - after Antimycin injection, 9) Non-mitochondrial respiration = after Rotenone / Antimycin injection. The assay was conducted with chambers heated to 37°C for a duration of approximately one hour. Results are expressed in pmol/(s*mL).

### Genomic DNA isolation of THC exposed sperm

DNA was isolated from THC exposed and unexposed sperm using a modified protocol from Quiagen Blood and tissue kit (60954). All centrifugation was conducted at 17,000 X G unless otherwise stated. Briefly, 3.5 mL Buffer 1 (NaCl 150 mM and EDTA10 mM) was added to sperm suspensions followed by vortexing for 10s at full speed and then centrifugation at 2500 X G for 5 min. The supernatant was removed to ∼ 1 mL and the sample vortexed again for 10s at full speed. Samples were then transferred to clean 2.0 mL microfuge tubes with the addition of Buffer 1 (0.5mL) back to the empty tube for an additional 10 sec vortex to collect residual sperm. The entire sample was combined and centrifuged for 2 min. The supernatant was discarded, and the pellet resuspend in 300 µL of Buffer 2 (TRIS-Cl 100 mM, EDTA,10 mM NaCL, 500 mM, SDS 1%).

Dithiothreitol (DTT, 1mM; 30µL) and 100uL of Proteinase K (1X) were added, and samples incubated at 55°C for 2 h with intermittent inversion. After incubation, 400µL of Buffer ATL was added and incubated for 10 min at 55^°^C. Samples were then passed through DNA binding columns by centrifugation for 1 min. Buffer AL + ETOH (400µL each) was added followed by centrifugation for 1 min and flow was discarded. Samples were placed in a new collection tube and 500µL of AW1 buffer was added followed by centrifugation for 1 min, and then 500 µL of AW2 buffer was added followed by centrifugation for 3 min. Final elution was accomplished with 50 µL RNase free water. Samples were stored at -20°C for subsequent analyses.

### Enzymatic methyl-seq conversion of genomic DNA isolated from sperm

Following DNA isolation, an Enzymatic Methyl seq Conversion kit module (NEB#E7125S) was used to convert gDNA into methy-converted DNA for PCR amplification and downstream epigenomic analyses in accordance with our optimized protocols [76]. Enzymatic Methyl seq conversion was achieved in a two-step process. The first step included a cocktail consisting of TET2 Reaction Buffer, Oxidation Supplement, DTT, Oxidation Enhancer, TET2, and Fe Solution added to 28 µL volumes containing DNA followed by incubating at 37°C for 1 h. The TET converted DNA samples were cleaned using AM Pure beads and final elutions were performed in 17uL of supplied buffer. In the second step, DNA was denatured with Formamide followed by deamination with APOBEC Reaction Buffer, BSA and APOBEC. Converted DNA was cleaned again using AM pure beads and final elution was achieved with 21uL of Elution Buffer. Converted DNA was then quantified using a Qubit DNA kit (ThermoFischer Cat No: Q32850) for further amplification and sequencing. The enzymatic methyl seq converted DNA from sperm were used for Whole genome Enzymatic Methyl Seq (EM-seq) analyses.

### Whole genome Enzymatic Methyl Seq *(EM-seq)*

Enzymatic detection of 5-methylcytosines (5mC) and 5-hydroxymethylcytosines (5hmC) was achieved with construction of libraries and sequencing by the Genomics Technology Core at the University of Missouri. Libraries were constructed following the manufacturer’s protocol with reagents supplied in the NEBNext® Enzymatic Methyl-seq Kit (New England Biolabs, Catalog #E7120). Briefly, fragmentation of gDNA isolated from THC exposed and unexposed sperm was performed on a M220 sonicator (Covaris). DNA fragments were end repaired/dA-tailed followed by adaptor ligation. A two-step enzymatic conversion protecting 5mC and 5hmC from deamination was performed prior to deamination of the cytosines residues. Fragments were PCR amplified to enrich libraries and incorporate barcodes. The final amplified cDNA constructs were purified by addition of Axyprep Mag PCR Clean-up beads (Axygen) selecting for an insert size of 550 bp. Libraries were quantified with the Qubit HS DNA kit (Invitrogen) and the fragment size analyzed on a 5200 Fragment Analyzer (Agilent). Libraries were diluted according to the standard protocol for sequencing on the NovaSeq 6000 (Illumina). The nf-core/methylseq (v2.6.0) automated bioinformatics pipeline [78] was utilized to analyze enzymatic methylation sequencing (EM-seq) data. This pipeline processes raw FastQ data, aligns reads, and conducts extensive quality control. The Bismark workflow within nf-core/methylseq was used to align reads to the *Bos taurus* genome (assembly ARS-UCD1.3) with Bismark [79]. The methylation coverage files (*.cov.gz) generated by Bismark, which include methylation percentages and read depth at each CpG site, were subsequently analyzed using the methyl Kit R package for downstream processing [80] By comparing the base conversion at cytosine sites between treated and untreated DNA, the methylation status (methylated vs. unmethylated) was determined. Differentially methylated regions between control and treatment (DMRs) were considered different with adjusted *q*-value of ≤0.05 and a ≥20% difference in methylation between THC exposed and VC samples.

### Functional Analyses of DMRs

Common DMRs revealed by EM-seq in THC exposed sperm compared to unexposed sperm were analyzed using Ingenuity Pathway Analysis (IPA, Qiagen) (n = 3 bulls, 2 replicates/treatment). Criteria for the set of common DMRs used in each analysis was determined based on an adjusted *q*-value of ≤0.05 with ≥3 CpG sites in sequenced promotor regions using multiple comparisons correction (i.e.,−log(Benjamini–Hochberg *P*-value) ≥ 1.3 or Benjamini–Hochberg *P*-value ≤0.05) that were predicted as increased or decreased based on a *Z*-score ≥ 2.0 and ≤ −2.0, respectively. The predictions were deemed to be activated or inhibited based on the *Z*-score criteria described above. IPA mapped the input genes to knowledge bases and identified the most relevant biological functions, networks, and canonical pathways related to the altered methylation profiles in the treatment and control. Expression analysis was designated as the core analysis type, and Expr Fold Change indicated as the measurement type [81–85]. The top canonical pathways associated with methylation difference are presented.

### Gene ontology and molecular function analysis

Gene ontology enrichment analysis was performed using the tool GO PANTHER version 19.0 (https://pantherdb.org/) to better understand the functional roles of DMR between THC exposed and unexposed control sperm. Genes were uploaded into the Protein Analysis through Evolutionary Relationships (PANTHER) classification system. The gene list analysis option was used to identify overrepresentation of gene ontology (GO) terms in the gene list data. The most significantly enriched ontologies were identified based on the list of molecular functions to which these genes were assigned [86–88].

### miRNA Quantification in Sperm

Total RNAs were isolated from frozen-thawed bovine sperm from both THC exposed and unexposed control after 24 h of exposure as described using Zymo Research Direct-zol RNA Mini Prep kit (Cat No: R2050S) according to the manufacturer protocol. All centrifugation was performed at 16000 x G for 1 minute. Briefly, 400 µL of sperm extended in TALP were added to 1200uL of TRIzol^®^ and centrifuged. Supernatant was transferred to equal volumes of ethanol (95-100%) and mixed thoroughly. The mixture was transferred into a spin column for centrifugation, and the column was transferred into a new collection tube followed by addition of 400 µl RNA wash buffer for centrifugation. DNase I (5µL) and DNA digestion buffer (35 µL) were mixed and applied to the same column matrix and incubated for 15 min at room temperature followed by addition of 400 µL of RNA prewash to the column for centrifugation. RNA Wash Buffer was added for centrifugation and RNA was eluted in 15 µL of DNase/RNase-Free Water Reverse transcription of total RNAs was accomplished using Qiagen miRCURY LNA^TM^ RT KIT (Cat NO: 339340) by a polyadenylated method according to the manufacturer’s protocol. Briefly, the reaction mixture consisted of 4ul 5X miRCURY Syber Green RT Reaction Buffer, 13uL of RNase-free water, 2uL 10X miRCURY RT ENZYME MIX and 1 uL of RNA template (20 µLtotal reaction volume). The thermocycler condition consisted of 60 min at 42°C, 5 min at 95°C, and 4°C hold.

The abundance of four miRNAs (bta-mir-449b, bta- mir-216a, bta-mir-502b and bta-mir-320) were quantified using real-time PCR. The miRNA sequences were derived from miRbase (https://www.mirbase.org/) and primers were designed using miRPimer [89] (Table. 1). Reactions were performed using Power SYBR Green PCR Master Mix in 12.5 µL volumes (Applied Biosystems, 4367659) and optimized primer concentrations (0.30 µM) in duplicate under the following conditions: 95°C for 10 min (initial denaturation), 40 cycles of 95°C for 15 s and 51°C for 1 min, followed by melt curve analysis of 65°C for 5 s and 95°C for 5 s. Duplicate no template control (NTC, water replacing cDNA) were included for each miRNAs in every run. The ^ΔΔ^Ct method was employed to determine and compare relative expression levels of each miRNA and was normalized against bta-mir-16 expressions. Two PCR replicates were performed for each treatment (VC and THC).

### Detection and quantification of DNA methyltransferase 1 (DNMT1) and DNA methyltransferase 3A (DNMT3A) in THC exposed sperm by immunoblotting

Characterization and quantification of methyltransferases (DNMT1/3A) were performed in THC exposed and unexposed sperm by western blot (WB) analyses. Protein extraction, immunoblotting, antibody incubation and imaging of the membrane was similar as mentioned above in detection and characterization of cannabinoid recptor-1 (CB1) section. Host species primary antibodies used for overnight incubation at 4C^0^ included Anti-DNMT1/3A (1:2000 dilution, GeneTex Cat No: GTX116011) and GTX129125, respectively). Densitometric analysis of protein bands was performed using the ImageJ software [90]. Relative expression levels of each enzyme were normalized against β-tubulin. Six replicates were performed for each protein of interest in both THC exposed and unexposed control.

### In vitro embryo production

Bovine ovaries were obtained from a local abattoir. Follicles ranging between 3 and 6mm in diameter were aspirated using a 20-gauge needle and syringe. Morphologically normal oocytes with a relatively uniform radial distribution of compact cumulus cells were selected and cultured for 24 h in Medium-199 with Earle salts (base medium) supplemented with 20 mM Na-pyruvate at 38.5°C and 5% CO2 in humidity saturated air. Frozen-thawed sperm from three bulls applied to 24 h THC 32nM exposure were then passed through a 45:90% Percoll step gradient consisting of a HEPES buffered Tyrode’s lactate sperm medium (99 mM NaCl, 24.8 mM NaHCO3, 10 mM HEPES, 0.33 mM NaH2PO4, 24 mM sodium lactate (60%), 2.4 mM MgCl2·2H20, 2.6 mM CaCl2·2H2O). Expanded cumulus-oocyte-complexes (COCs) were co-incubated with pooled motile sperm at a ratio of 8000:1 sperm per oocyte for ∼15 h in fertilization medium (114 mM NaCl, 25 mM NaHCO3, 3.2 mM KCl, 0.39 mM NaH2PO4, 0.18 mM penicillin-G, 16.6 mM sodium lactate, 0.76 mM MgCl2·2H2O, 2.7 mM CaCl2·2H2O) at 38.5°Cin 5% CO2 and humidity saturated air [91, 92]. Presumptive zygotes were stripped of cumulus by vortexing and selected for incubation in KSOM culture medium (EMD Millipore, Billerica, MA) supplemented with 0.3% BSA at 38.5°C and 5% CO2 in atmospheric oxygen. No more than 20% of presumptive zygotes were discarded from culture following fertilization to ensure unbiased selection among embryos fertilized from THC exposed and unexposed sperm. The culture medium was supplemented with 5% FBS at 72 h post-fertilization. Embryos were evaluated for cleavage at 48 h (D2), 8–16 cell stage at 72 h (D3) and for blastocyst development at 180 h (D7.5). N = 5 separate replicates (sperm from 2 bulls combined in each replicate) were performed for both THC and VC.

### Statistical Analysis

All data were analyzed by logistic regression using a generalized linear mixed effect model with SAS 9.4 software (St. Louis, Missouri, USA). The experimental unit was the sperm samples to which the treatment was applied and therefore considered as a fixed effect. Sperm kinematic, Oroboros and TUNEL data were analyzed by a mixed model in which treatment, time, and treatment *time were considered as fixed effects in the model while bull and replicate were assigned as random effects. Because sperm samples from bulls were pooled for developmental and flow cytometry data, bull effect was excluded from the model. Exposure (THC or VC) were assigned as fixed effects and replicates were random variables. Treatment*time interaction was also included for flow data as a fixed effect as the experiment was conducted both at 0 and 24 h. For determining the relative expression of miRNAs and DNMTs, a mixed model was used with treatment and bull as a fixed effect and replicate as a random effect. Residuals for all parameters were evaluated for normal distribution (Shapiro-Wilk test). Log (base) transformations were applied where normality distributions were not met (P≤0.05). Where data did not meet the normality distribution, LSM and SE were presented as the back transformed value.

## Results

### Localization and characterization of Cannabinoid receptor-1(CB1)

Cannabinoid receptor-1 (CB1) was detected in post-ejaculatory bovine sperm maintained in non-capacitating conditions at the post-acrosomal region by immunofluorescent staining of fixed sperm. Human placenta, bovine placenta and murine brain served as positive controls for expression of CB1 **(Fig. 1A)**. A single and specific band was detected by western blot at the expected molecular weight of **∼**75kD using equine pituitary as a positive control and thus further authenticating CB1 detection in mature bovine sperm **(Fig. 1B)**. Sperm were then held in both non-capacitating or capacitating conditions for 6 hr to determine changes to CB1 detection. CB1 detection was decreased in capacitated vs non-capacitated samples as determined by immunofluorescence staining (**Fig. 2)** Sperm challenged by THC exposure in non-capacitating conditions displayed loss of acrosome integrity and attenuated CB1 expression despite remaining uncapacitated, as shown by lack of tyrosine phosphorylation.

**Fig 1.**
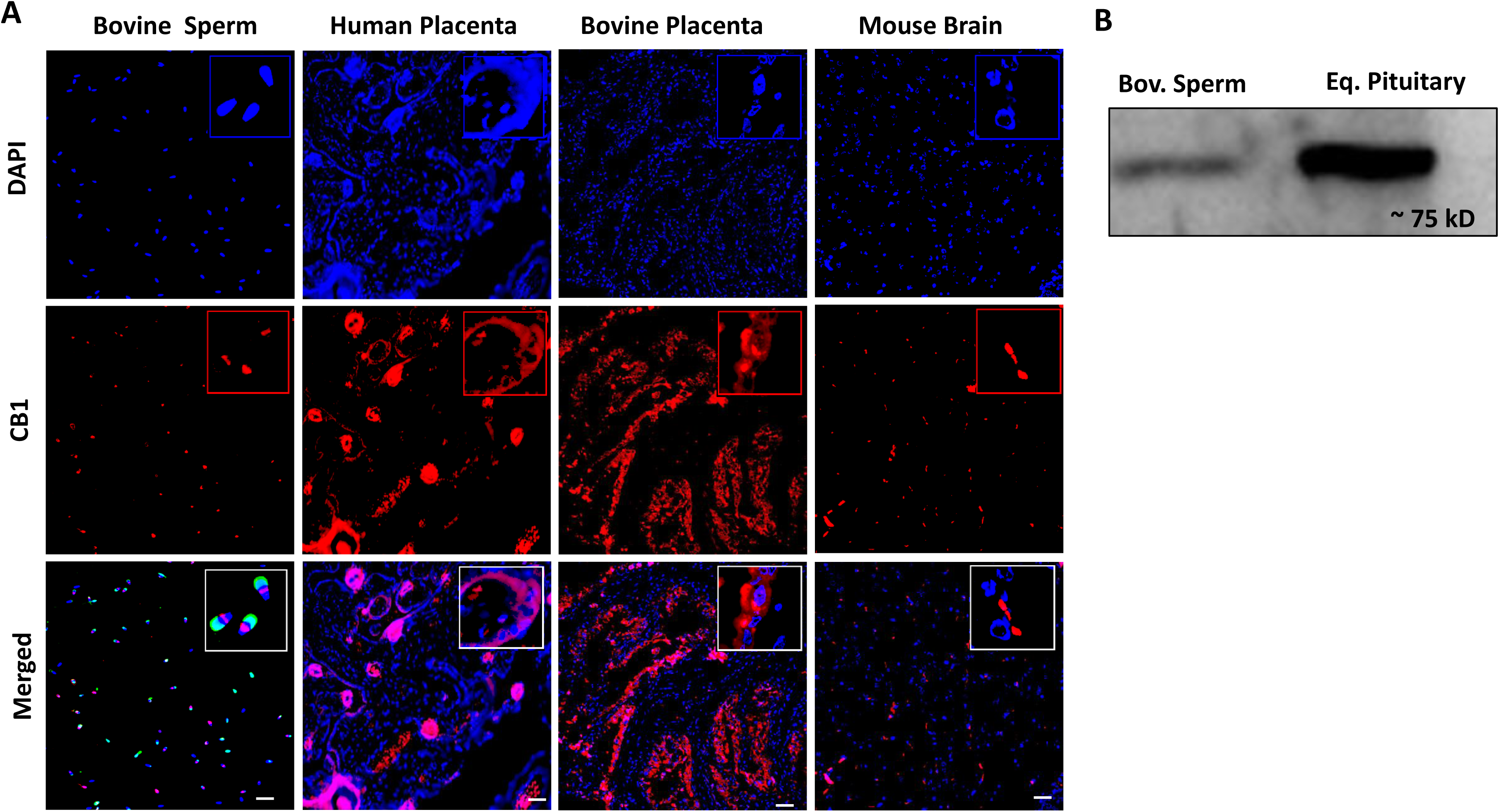
Localization and characterization of the CB1 receptor in sperm. A) Immunocytochemistry images of bovine sperm, human placenta, bovine placenta, and mouse brain depicting nuclear staining with DAPI (blue), CB1 (red), and merged images. FITC-PSA fluorescence can be observed in the merged image of sperm (green), for identification of the sperm acrosome. B) Western blot image of bovine sperm and equine pituitary identifying a single and specific band at the expected 75 kD value. Scale bar = 50 μM.

**Fig 2.**
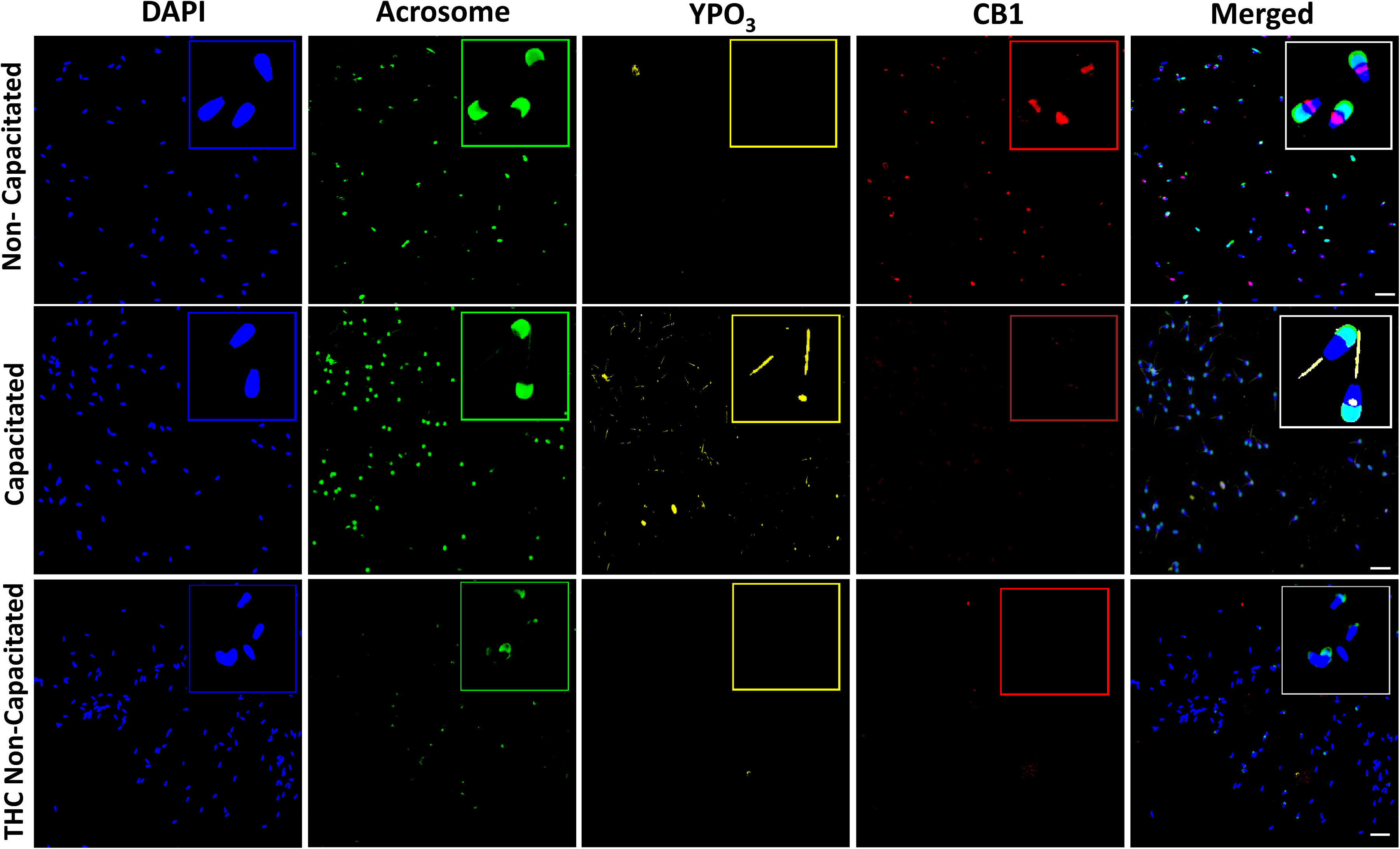
Influence of THC exposure and capacitation status on CB1 detection in sperm. Localization of CB1 in non-capacitated sperm (upper panel), to the post-acrosomal sheath as compared to loss of abundance when sperm are held in capacitating conditions (middle panel) as indicted by tyrosine phosphorylation staining (YPO_3-,_ yellow). Sperm exposed to THC under non-capacitating conditions exhibit decreased YPO_3_, CB1 and acrosome detection (green) (lower panel). Scale bar = 50 μm.

### THC compromises acrosome integrity in sperm

Postejaculatory sperm were subjected to flow cytometry for detection and quantification of acrosome integrity (FITC-PSA) and capacitation status (merocyanine) following THC exposure for 24 h (**Fig. 3A-D**). THC exposure to sperm did not have an immediate affect on sperm viability or acrosome integrity **(Supp. Fig 1)**. However, sperm exposed to THC for 24 h experienced a significant loss in acrosome integrity without affecting viability (67% vs. 2%, respectively; **Fig. 3A,C**). Similarly, the capacitation status of THC exposed postejaculatory sperm was also evaluated at 0 and 24 h. Unlike acrosome integrity, exposure of sperm to THC for 24 h did not alter the capacitation status of sperm (*P*=0.7) (**Fig. 3B,D**).

**Fig 3.**
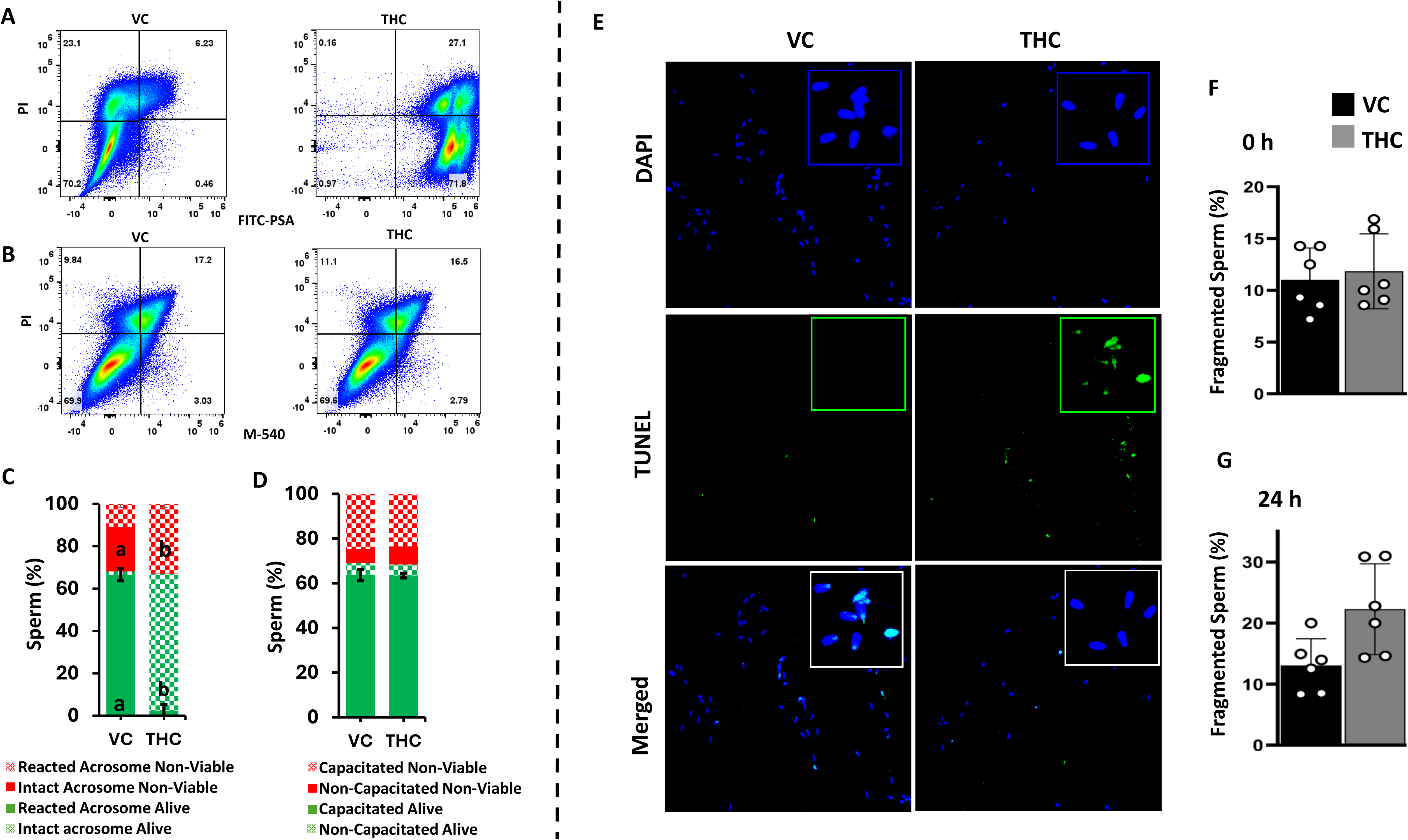
THC exposure to sperm compromises acrosome and DNA integrity. Postejaculatory sperm were exposed to THC for 24 h and subjected to flow cytometric analysis for determination of acrosome integrity by FITC-PSA plotted against viability using propidium iodide staining (PI) as a marker of membrane permeability (**A, C**). Sperm were also quantified for THC effects on capacitation status by Merocyanine (M540) staining and plotted against PI fluorescence (**B,D**). THC exposure caused an increase in acrosome reacted sperm (**C**) without affecting capacitation status or viability (**D**). Sperm were also subjected to a DNA fragmentation assay by TUNEL following exposure to THC for 24 h (**E**). Nuclear staining was detected by DAPI (blue), while DNA fragmentation was indicated by FITC (green), and categorized as fragmented and not fragmented after initial exposure (**F**) and again after 24 h (**G**). n = 3 bulls; 2 individual replicates/bull. Scale bar = 50 μM. Different superscripts (^a,b^,) indicate P<0.05 between treatments.

### THC exposure to post-ejaculatory sperm increases DNA fragmentation

The effects of THC exposure on the DNA integrity of post-ejaculatory sperm were quantified by TUNEL assay at 0 and 24 h following addition of THC to sperm under non-capacitating conditions. DNA fragmentation was categorized as fragmented or not fragmented **(Fig. 3 E-G)**. THC exposure had no effect on DNA fragmentation at 0 h **(Fig. 3F)**, but THC exposure to sperm for 24 h elicited loss of DNA integrity (*P=*0.03). The percentage of DNA fragmented sperm was higher in THC group (22) compared to VC (13) at 24 h of exposure (**Fig. 3E,G**).

### THC exposure lowers progressive motility but does not alter direct mitochondrial bioenergetics potential of postejaculatory sperm

The effects of THC exposure on sperm motility kinematics as measure of indirect mitochondrial function were quantified by CASA at both 0 and 24 h of exposure (**Fig. 4A**). Sperm kinematics parameters included total motility (TM), progressive motility (PM), average path velocity (VAP), and curvilinear velocity (VCL). THC exposure did not alter motility parameters of sperm except for progressive motility, which was significantly but moderately lower (29% vs 36%, respectively) compared to unexposed sperm (**Fig 4A**). Similarly, there was a trend for THC to lower VCL of sperm after 24 h (106 vs 188, respectively; P=0.08, **Fig. 4A**). Mitochondrial respirometry traits from bull sperm exposed to THC and unexposed sperm held in suspension were quantified at two time points (0 and 24h) using the O2k system to measure direct effects on mitochondrial function **(Fig. 4B)**. Basal respiration decreased over time regardless of treatment. There was no significant effect of THC exposure on any sperm bioenergetic potential parameters (*P*>0.05) **(Fig. 4B)**, except for a trend of higher ADP and ATP linked and spare respiration in THC exposed sperm as compared to unexposed controls (*P*=0.06 - 0.1) **(Fig. 4B).** However, some of the respiration parameters which were measured decreased over time independent of exposure, indicating a significant time effect **(Supp Fig 2).**

**Fig 4.**
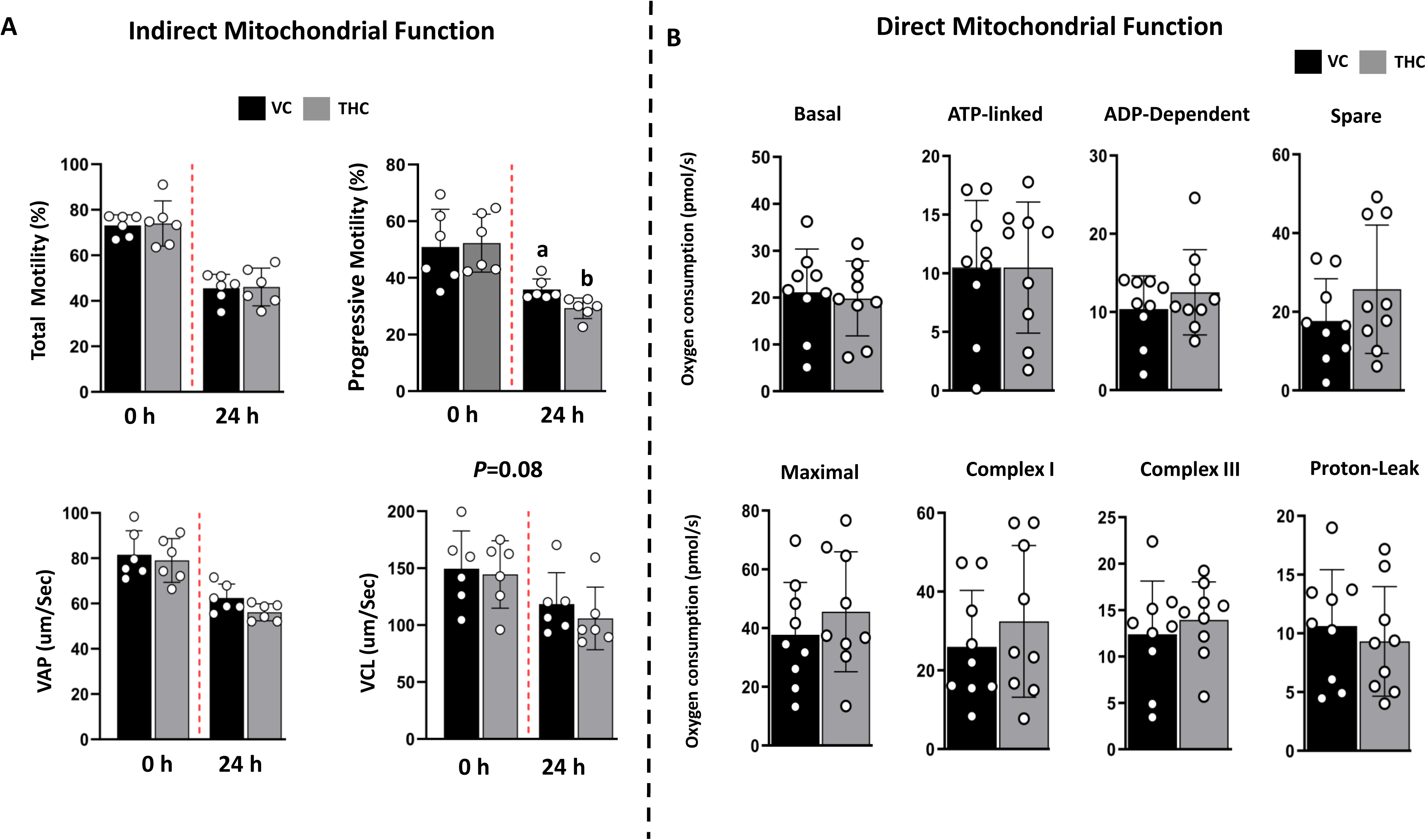
Effects of THC exposure on indirect and direct mitochondrial bioenergetics of sperm. Sperm were exposed to THC for up to 24 h at ambient temperature in non-capacitating conditions and indirect effects on mitochondrial function were determine by computer-assisted sperm analyses (**A**) for quantification of total and progressive motility average-path velocity (VAP), and curvilinear velocity (VCL) at 0 and 24 h. Direct assessment of mitochondrial function was conducted using high-resolution sperm respirometry (O2K) for quantification of respirometry values (**B**) including: Basal, ATP-linked, ADP-dependent, Spare, Maximal, Complex I, Complex III and Proton Leak respiration. Data are presented as mean ± SEM. n = 3 bulls; 3 individual replicates/bull. Different superscripts (^a,b^) indicate P<0.05 between treatments.

### THC exposure alters the miRNA abundance in sperm and compromises in vitro embryo development

Sperm exposed to THC were characterized for changes to the relative abundance of four miRNAs (miR-320, miR-520, miR-449, and miR-216) established for roles in embryo development and epigenetic programming (**Fig 5**). The relative abundance of miR-449 was lower in exposed versus unexposed control (*P=*0.04) while bull (*P=*0.0007) and bull*treatment (*P=*0.002) also had significant effects. Similarly, the relative abundance of miR-216 was lower (*P=*0.02) in exposed versus unexposed control while bull (*P=*0.30) and bull*treatment (*P=*0.63) have no significant effect. Contrary to that the relative expressions of miR-320 (P=0.8950) and miR-520 P=0.6586) were not altered between two treatment groups (**Fig 5B**).

**Fig 5:**
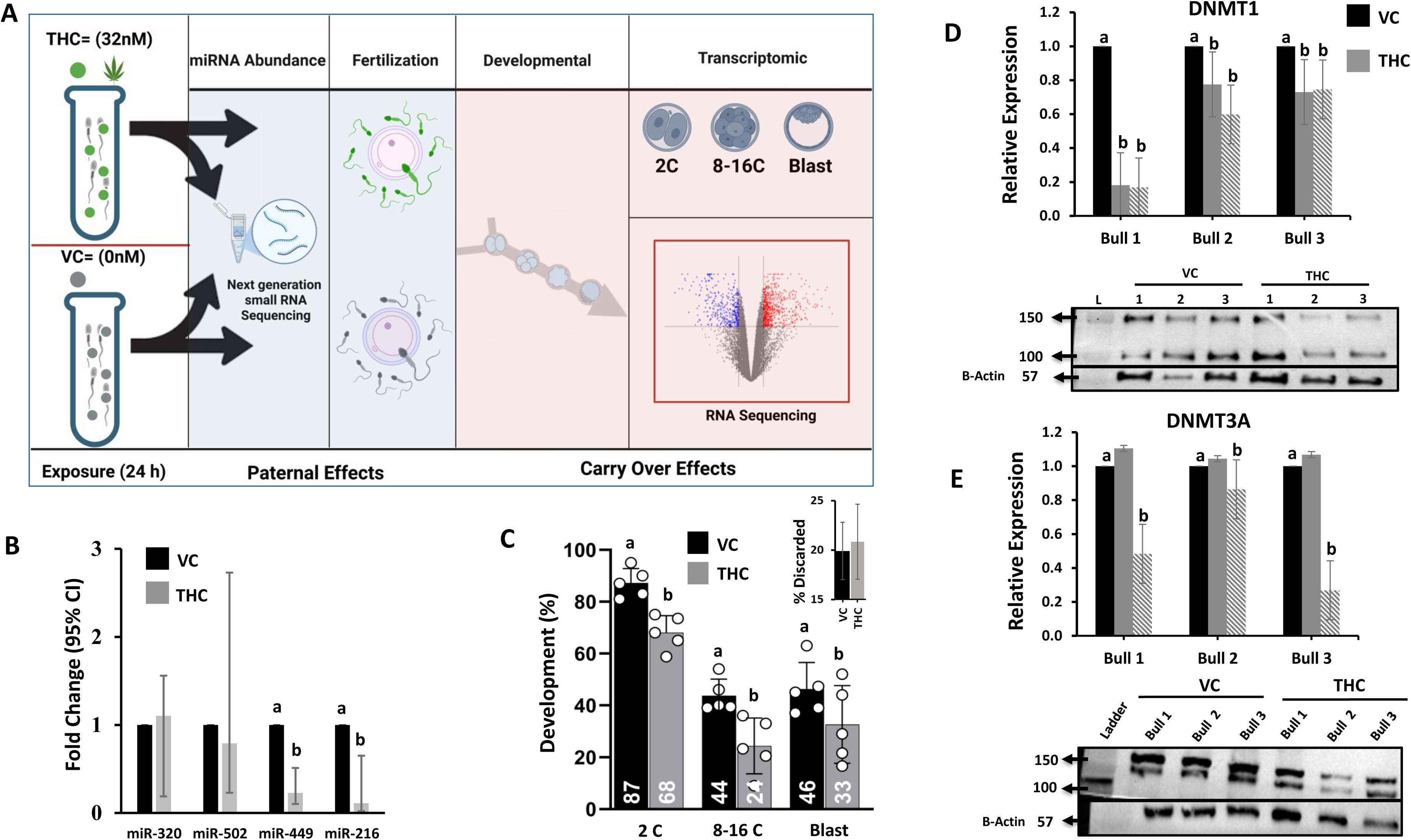
THC exposure to postejaculatory sperm reveals new epigenetic mechanisms with paternal carryover effects that decrease embryo developmental competence. Experimental model depicting postejaculatory sperm exposure to THC applied to an in vitro embryo continuum model for determination of paternal carryover effects (A). The relative abundance of four miRNAs was quantified and presented as foldchange against housing keeping miRNA XX? normalized to 1 (B). Pooled frozen-thawed bull sperm (n = 2) were exposed to THC for 24 h in non-capacitating medium, washed by centrifugation and applied to gamete co-incubation. The percent of two-cell embryo cleavage (2 C), 8–16 cell (8-16 C), and blastocyst development were reduced in embryos fertilized from THC-exposed sperm (C). The relative protein abundance of DNMT1 (D) and DNMT3A (E) including two splice variants, respectively were quantified in THC-exposed compared to unexposed sperm for individual male by western blot. Different superscripts (^a,b^) indicate P<0.05 between treatments.

Deficiencies of frozen-thawed bovine sperm function following exposure to THC were further investigated by determining incompatibility for vitro embryo production through blastocyst development. Sperm were exposed to THC for 24 h as previously described, washed twice by centrifugation and adjusted for equal concentration by motility compared to unexposed (VC) sperm for in vitro fertilization (1 × 10^6^ motile sperm/mL). The percent of embryo two-cell cleavage (68% vs 87% *P*=0.0133), 8–16 cell morula development (24% vs 44% *P*=0.0082) and blastocyst formation (33% vs 46% *P*=0.04) were reduced in embryos fertilized by THC-exposed sperm compared to unexposed sperm, respectively The percentage of embryos discarded after fertilization and prior to cleavage was not different (20% vs 21%) from embryos produced with THC-exposed versus unexposed sperm **(Fig. 5C**).

### THC exposure lowers the relative abundance of DNMT3A/1

The relative abundance of DNMT1/3A were quantified following sperm exposure to THC for 24 h. Two splice/transcript variants were detected in postejaculatory sperm for both DNMT1 and 3A. Variants identified for DNMT3A included independent bands at ∼130 kDa and 110 kDa, respectively. Similarly, detection of two variants for DNMT1 included 150 kDa (variant 1) and 100 kDa (variant 2). THC exposure to sperm reduced expression of both DNMT1 variants compared to unexposed sperm irrespective of bulls **(Fig 5D)**. THC exposure resulted in lower abundance of DNMT3A/2 in all three males compared to unexposed sperm. No significant effect of THC exposure was detected for variant 1 of DNMT3A/1 (**Fig. 5E**). DNMT1/3A variant detection in sperm was evaluated across all three independent bulls following THC exposure. Statistical modeling that included bull as a fixed effect revealed a significant effect of bull on DNMT 1/3A variant levels, indicating inter-individual variability. Although expression of DNMT3A/1 and both variants of DNMT1 was altered in response to THC exposure in all three bulls, the magnitude of this response differed among individuals. These results demonstrate that while THC exposure is associated with changes in DNMT3A variants, individual bull-specific differences contribute to variability in the observed response **(Fig 5D,E)**.

### THC exposure alters the methylation of genome-wide promoter regions in mature sperm

Principal component analyses (PCA) applied to whole-genome EM-seq promoter sequences of sperm revealed distinct clusters unique to individual males regardless of THC exposure **(Fig. 6A)**. Comparison of sperm exposed to THC *versus* VC revealed ∼56 DMRs, 39 of which were hypomethylated compared to 17% hypermethylated irrespective of male effect (**Fig. 6B)**. THC exposure induced both male-specific and shared alterations in sperm DNA methylation across the three bulls, as illustrated by the Venn diagram of differentially methylated regions (DMRs). Each bull exhibited a distinct set of unique DMRs (Bull A: 17; Bull B: 30; Bull C: 21), indicating variability in the epigenetic response to THC exposure among individuals. Moreover, 3 shared DMRs were observed between Bull A and B and 1 in each between Bull B and C and Bull A and C, as well as a subset of DMRs (n = 3) common to all three bulls **(Fig 6B)**. Gene ontology and molecular function enrichment analysis using GO Panther indicated that common identifiable DMRs were mainly related to the functional categories of binding (22%) and catalytic activity (15%), while the largest proportion (48%) were not assigned due to lack of annotation **(Fig. 6C)**.

**Figure 6:**
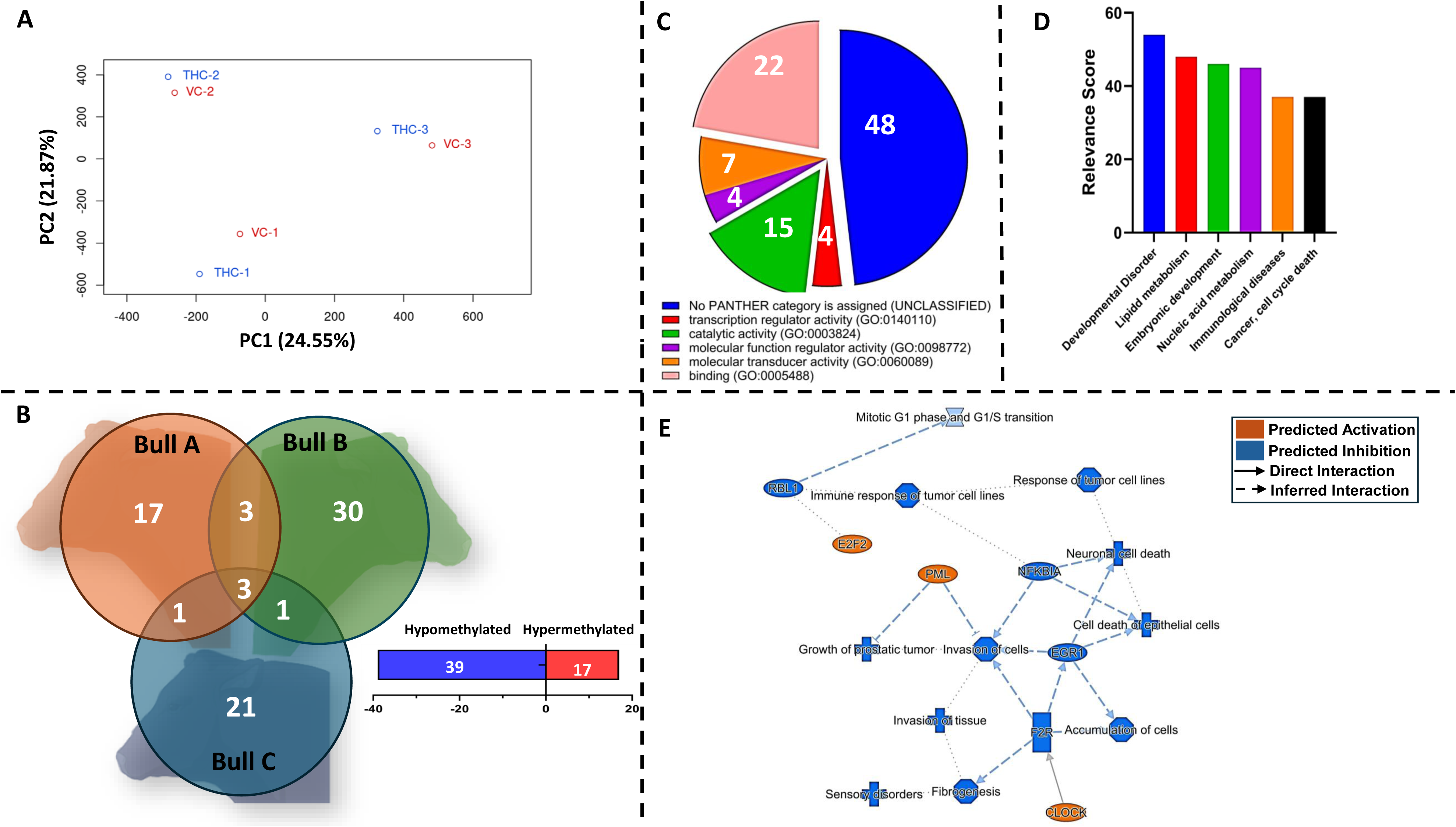
Genome-wide methylation status and impacted biological functions resultant of postejaculatory sperm exposure to THC. Whole-genome Em-seq was conducted to identify changes to the promoter regions of postejaculatory sperm exposed to THC for 24 h compared to vehicle control (VC − 0.1% DMSO). Principal component analysis of THC and VC-exposed sperm in which all three groups form distinct clusters (**A**). The percent variance explained by PC1 and PC2 is 25% and 22%, respectively. Venn diagram of differentially methylated regions between three males and total numbers of DMRs between THC vs. VC irrespective of male (**B**). Molecular function and gene ontology of THC-affected DMRs as identified by PANTER included binding (22%), catalytic activity (15%), and unclassified categories (48%) (**C**). Characterization of bio-function categories by relevance and significance score where canonical pathways and function annotations were classified (**D**). Graphical summary of IPA for DMRs between THC and VC groups indicated that Mitotic G1 phase is the major pathway predicted to be inhibited by THC leading to inhibition of response of tumor cell lines, immune response and neuronal cell death and cell death of epithelial cells (**E**). Activation or inhibition *Z*-score of the top 15 most significant canonical pathways affected by THC. Orange indicates activated pathway, blue is inhibited, white means no activity, and gray indicates that no activity pattern could be measured **(F**). *n* = 3 bulls; 2 pooled replicates/treatment.

### Functional analyses of DMRs associated with THC sperm exposure include the pathways of developmental disorder, embryo development and mitotic G1 phase

IPA analyses identified 6 bio-function and disease categories based on relevance score as a result of THC exposure altering DMRs within the promoter regions of mature sperm. The major categories which were identified included developmental disorder, lipid metabolism, embryonic development, nucleic acid metabolism, immunological diseases, and cancer and cell death **(Fig. 6D)**. A network analyses of affected DMRs indicated that Mitotic G1 phase is the major pathway predicted to be inhibited by THC and is associated with inhibition of immune response of tumor cell lines and NFKBIA pathway, leading to cell death of epithelial cells (**Fig. 6E**). Canonical pathway analysis with common DMRs between THC exposure and control that were mapped by IPA led to 65 significant pathways. The top 15 most significant pathways presented were mostly related to mitotic G1 phase/G1 transition, Huntington disease signaling, synthesis of DNA and S phase senesce pathways **(Supp Fig. 3).**

## Discussion

Postejaculatory spermatozoa are a highly specialized terminal cell that must transit through a dynamic environment to deliver intracellular contents for successful fertilization. Transport through the female reproductive tract influences sperm behavior by altering mitochondrial respiration, motility and capacitation events for gamete interaction. Despite conventional dogma suggesting that paternal carry over effects are limited to events during spermatogenesis, we provide new evidence of THC exposure that specifically targets previously unestablished genomic and epigenomic mechanisms. Although increased cannabis use certainly justifies an urgent need to determine direct reproductive consequences of THC on mature sperm exposure, these newly defined mechanisms and carryover effects emphasize the overlooked consequences of environmental perturbations to postejaculatory sperm independent of spermatogenesis.

The experiments herein included 24 h THC exposure to mature postejaculatory sperm to mimic temporal aspects of sperm transit after semen deposition in the female reproductive tract. The detection and characterization of CB1 expression provides mechanistic evidence for THC interaction with mature sperm. Receptor-mediated CB1-THC interaction is well-established in the central nervous system, where THC acts as a partial agonist to elicit psychoactive effects [93, 94]. Downstream intracellular responses of THC-CB1 interaction include increased intracellular calcium, adenylate cyclase inhibition and reduced cAMP [94–96], thereby inhibiting neurotransmission [94, 97–99]. In agreement with these data, mature sperm exposure to THC resulted in no acute changes to motility kinematics and only a slight reduction in progressive motility. Similarly, no direct effects on mitochondrial function were detected when analyzing multiple indices of mitochondrial bioenergetics using high-resolution sperm respirometry. These data also agree with failure of THC exposure to elicit capacitation changes to sperm as quantified by flow cytometric values. The agonistic effects of THC on the CB1 receptor within the endogenous cannabinoid system align with known mechanisms whereby mature sperm undergo capacitation, which are dependent upon a rise in intracellular cAMP and calcium to stimulate sperm hyperactivation and phosphorylation of tyrosine residues (cite). Although not directly challenged in capacitating conditions, these results suggest that THC interacts with CB1 through canonical mechanisms that do not alone facilitate capacitation of sperm. However, THC exposure to sperm resulted in a complete shift from acrosome intact to acrosome-compromised sperm, suggesting that exposure to THC in seminal fluid or the female reproductive tract may be sufficient to elicit precocious structural changes that likely impact sperm penetration through the zona pellucida. Indeed, the functional status of sperm was partially challenged within an in vitro embryo production system, where we observed a reduction in 2-cell cleavage and subsequent embryo development, which is indicative of reduced sperm function. Similarly, previous experiments conducted by our group determined that exposure of sperm to the organotin tributyltin chloride was sufficient to cause acrosome degradation and also significantly reduced embryo cleavage and prevented blastocyst development [74, 100]. Loss of acrosome integrity is physiologically preceded by capacitation that is initiated within the oviduct (cite). The impact of THC on acrosome integrity but not capacitation appears autonomous and thus likely due to direct impacts on structural function rather than classical activation of pathways associated with capacitation due to no observable changes in merocyanine by flow cytometry. Collectively, these findings may prove beneficial for identifying appropriate assisted reproductive procedures for human clinical applications such as intracytoplasmic sperm injection (ICSI) that bypass the requirement for gamete interaction and thus can be applied to overcome structural defects such as loss of acrosome integrity that hinder sperm penetration during gamete interaction.

Although current dogma suggests that mature sperm chromatin is largely protected from external insult due to protamine compaction, ∼10% retain euchromatin status and thus are susceptible to environmental insult [100]. An additional attribute of recreational drugs and environmental toxins is the demonstrated ability to interact and bind DNA albeit with unknown consequences on downstream effects following fertilization and subsequent embryo development [100–105]. THC exposure to sperm for 24 h elicited significant reductions in DNA integrity. The lack of overt phenotypical characteristics but compromised genomic integrity of THC-exposed sperm suggests that paternal consequences may be idiopathic simply due to lack of routine clinical diagnostics that fail to identify intracellular mechanisms impacted by THC exposure. Concerningly, such changes may manifest in the embryo as carryover effects originating from compromised nuclear function of sperm. Indeed, recent traction on the importance of the sperm epigenome has provided valuable insight into the functions of miRNAs and other non-coding RNA- associated mechanisms important for post-fertilization events [81–85]. The sperm epigenome plays a crucial role in many biological processes including early embryo development, zygotic genome activation, imprinted clusters, and X chromosome inactivation [106–110]. Environmental factors that alter DNA methylation, non-coding RNAs, and histone modifications may impact embryo development with transgenerational effects [111–116]. Environmental factors that induce epigenetic modifications to mammalian sperm have been described during spermatogenesis [117–126], but studies to determine potential effects on postejaculatory sperm have not been described. The present study utilized a comprehensive genome-wide methylation analysis of postejaculatory bovine sperm to determine effects of THC on the promoter regions of the sperm methylome. Interestingly, distinct clustering among three different males suggests individual susceptibility to THC exposure that may be explained by a male effect, heterogeneity of sperm within a sample, or diversity of chromatin accessibility within a population. However, whole EM-Seq promotor regions included ∼ 56 DMRs as a result of direct THC exposure, irrespective of the individual male. In agreement with these data demonstrating environmental impacts to the postejaculatory sperm methylome, our earlier study determined that TBT exposure resulted in 750 DMRs in mature sperm [100]. IPA analysis of THC-exposed sperm identified DMRs associated with developmental disorder, lipid metabolism, mammalian embryo development and nucleic acid metabolism, cancer, and cell death. More specifically, inhibition of mitotic G1 phase and G1/S transition was indicated as the most likely affected pathways, which may seem inconsequential in the terminally differentiated sperm while contrastingly concerning for post-zygotic embryo cleavage requirements following fertilization.

Such genome wide methylation changes in mature sperm are unprecedented because canonical mechanisms indicate a requirement for de novo methyltransferase activity, whereas postejaculatory sperm are both transcriptionally and translationally quiescent. Although mammalian sperm do indeed possess DNMTs and TETs as remnants of spermatogenesis [127–130], their functionality in mature sperm was not previously determined. The current data set provides strong evidence for DNMT1/3A in mature sperm with a decrease in relative abundance of splice variants in response to THC exposure. Such observations provide a new mechanism for alterations to the methylome of mature sperm that are independent of transcription.

In support of these newly proposed mechanisms that affect the methylome of mature sperm, roles of miRNA delivery from sperm to the embryo have recently been established [131–133]. However, direct consequences of environmental exposure to miRNA level abundance in mature sperm have not previously been explored. When exposed to THC, both miR-216 and 449 abundance was decreased. These miRNAs are primary drivers of cell cleavage, cell proliferation and epigenetic reprogramming of early embryos [134, 135]. In agreement with decreased levels of miR-216 and 449 in sperm and known function in early embryo development, groups of embryos derived from THC-exposed sperm had significantly decreased development rates up to the blastocyst stage. Collectively, these findings suggest a direct paternal carryover effect to the early embryo and substantiate a need to define causal relationship.

## Conclusion

Results from experiments herein introduce a paradigm-shift with proposed mechanisms to advance understanding of environmental perturbations to mature sperm that confer epigenetic alterations. Further studies will determine putative carryover methylation changes of paternal origin and their impacts on early embryo development.

## Ethics Declaration

The authors declare no conflict of interest that could be perceived as prejudicing the impartiality of the research reported.

## Contributions

MSS was responsible for data collection, analyses, and preparation of the manuscript. SA performed NGS data analysis and manuscript edits. JDAL, and RKB made significant contributions to data analysis, interpretation, and manuscript review. ZJ contributed to epigenetic concepts. BD responsible for experimental design, theory, analyses, manuscript preparation, and original concepts.

---

**Table 1.**
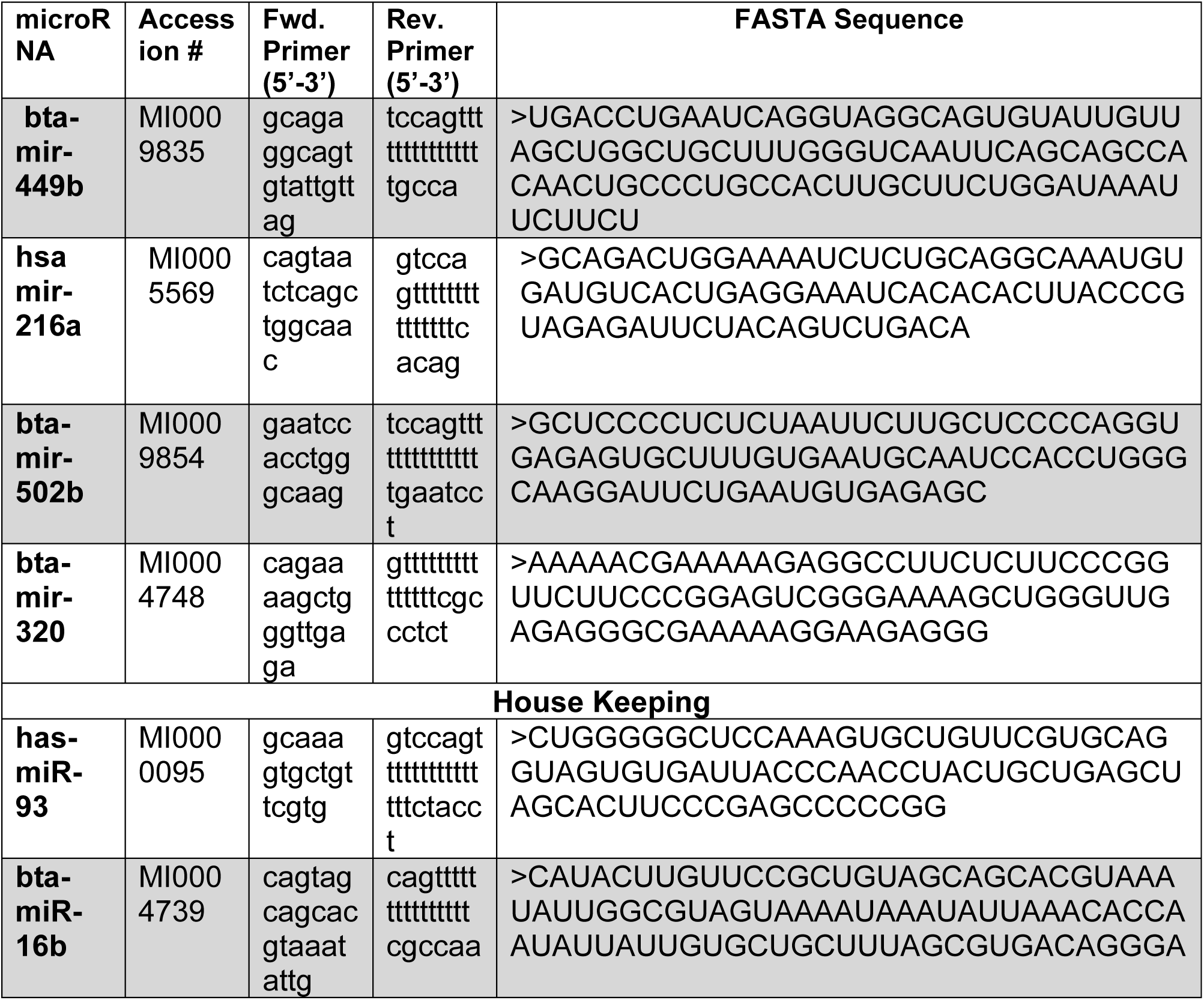
miRNAs primer sequences and accession numbers.

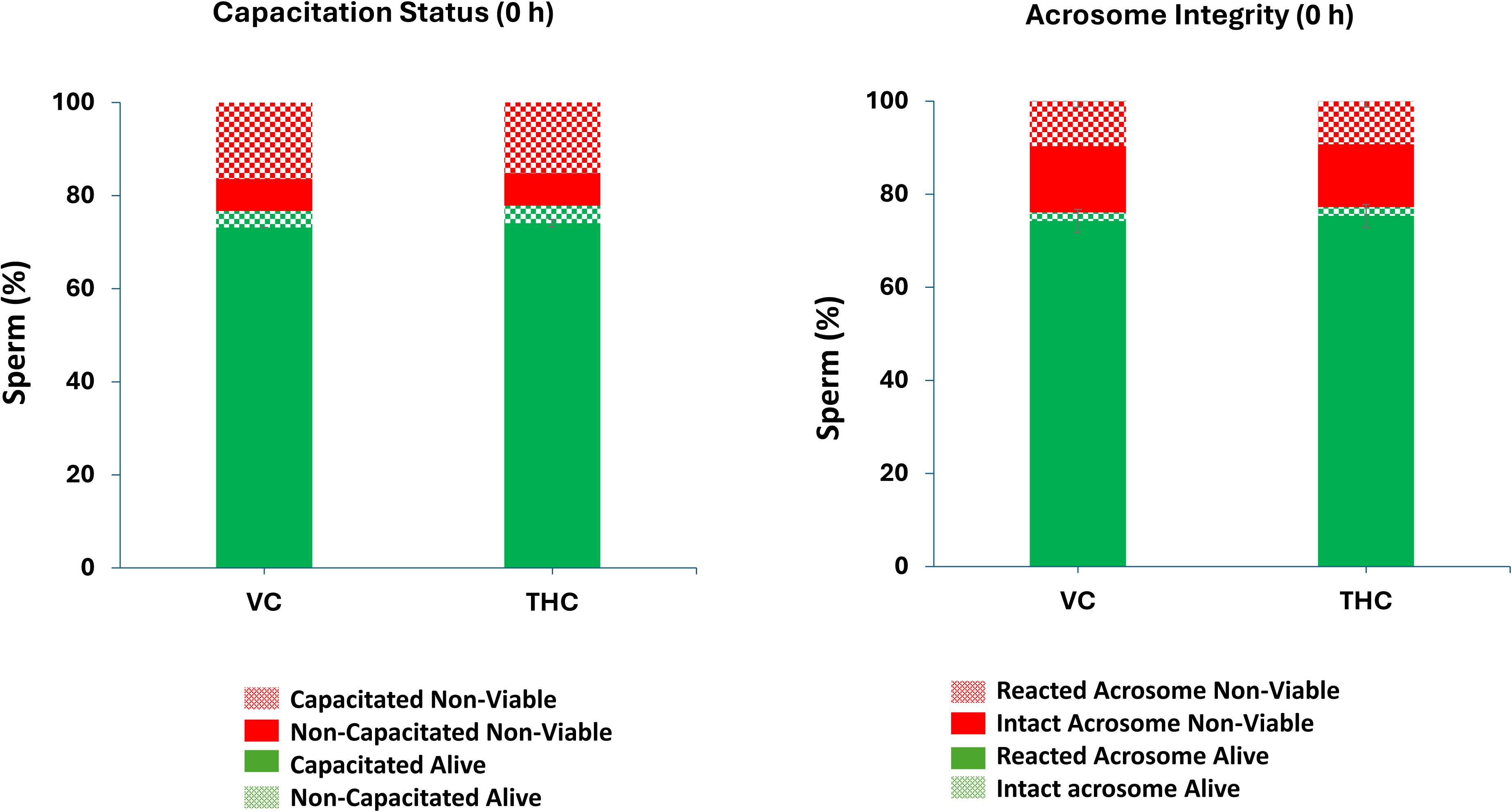

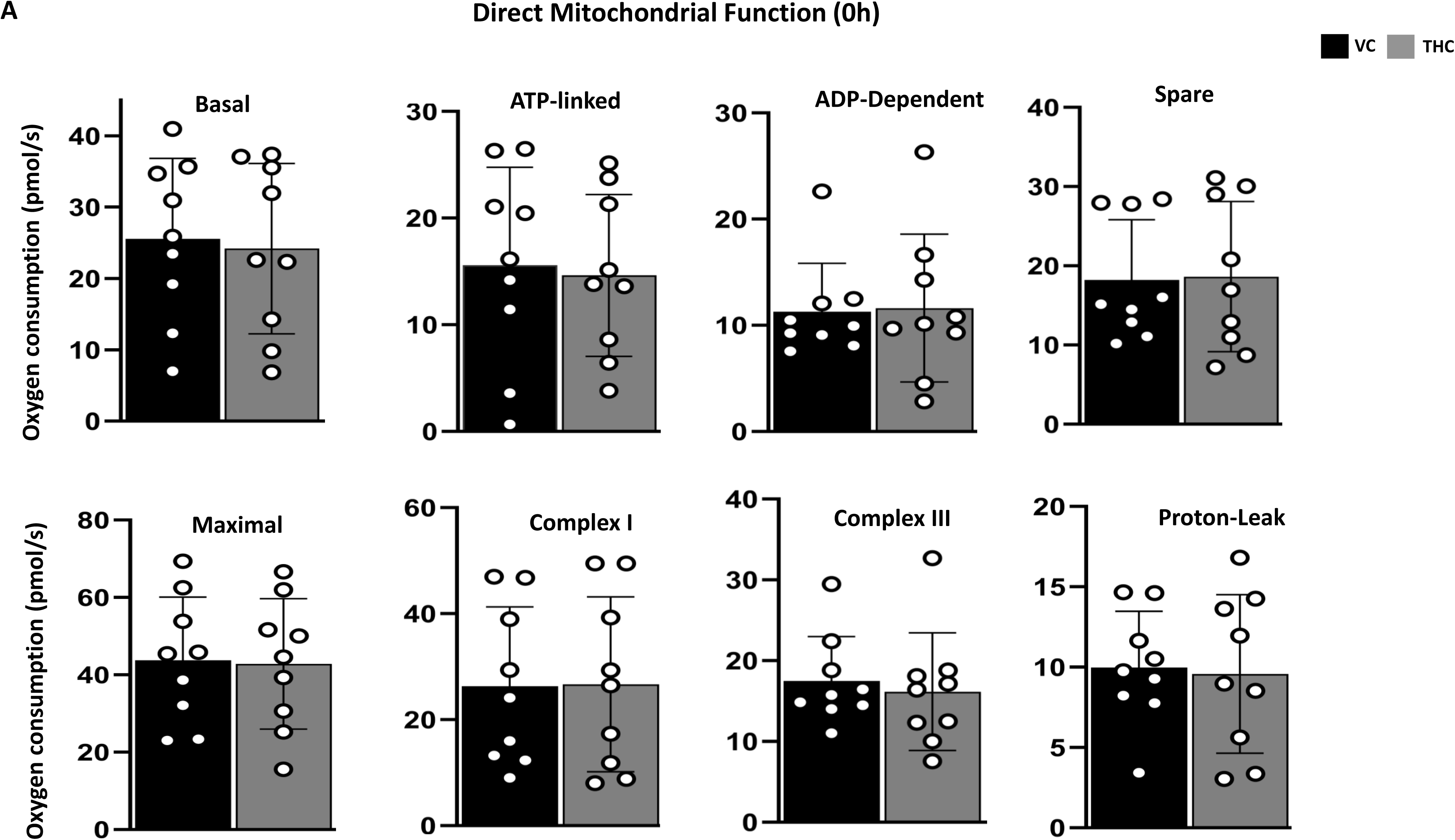

